# Clinically applicable rapid susceptibility testing of multi-drug resistant *Staphylococcus aureus* by mass spectrometry and extreme gradient boosting machine

**DOI:** 10.1101/2021.10.05.463151

**Authors:** Zhuo Wang, Hsin-Yao Wang, Yuxuan Pang, Chia-Ru Chung, Jorng-Tzong Horng, Jang-Jih Lu, Tzong-Yi Lee

**Author notes:** These authors contributed equally to this work (Zhuo Wang, Hsin-Yao Wang, Yuxuan Pang). To whom correspondence should be addressed: **Tzong-Yi Lee**, Warshel Institute for Computational Biology, School of Life and Health Sciences, The Chinese University of Hong Kong, Shenzhen, 2001 Longxiang Road, Longgang District, Shenzhen, 51872, China.; **Jang-Jih Lu**, Department of Laboratory Medicine, Chang Gung Memorial Hospital at Linkou, Taoyuan City, Taiwan.

## Abstract

Multi-drug resistant *Staphylococcus aureus* is one of the major causes of severe infections. Due to the delays of conventional antibiotic susceptibility test (AST), most cases were prescribed by experience with a lower recovery rate. Linking a 7-year study of over 20,000 *Staphylococcus aureus* infected patients, we incorporated mass spectrometry and machine learning technology to predict the susceptibilities of patients for 4 different antibiotics that can enable early antibiotic decisions. The predictive models were externally validated in an independent patient cohort, resulting in an area under the receiver operating characteristic curve of 0.94, 0.90, 0.86, 0.91 and an area under the precision-recall curve of 0.93, 0.87, 0.87, 0.81 for oxacillin (OXA), clindamycin (CLI), erythromycin (ERY) and trimethoprim-sulfamethoxazole (SXT), respectively. Moreover, our pipeline provides AST 24–36 h faster than standard workflows, reduction of inappropriate antibiotic usage with preclinical prediction, and demonstrates the potential of combining mass spectrometry with machine learning (ML) to assist early and accurate prescription. Therapies to individual patients could be tailored in the process of precision medicine.

## Introduction

Early detection of drug resistance of bacteria in patients is critical to prevent the spread of some pathogens in the epidemiology of infectious diseases. The extensive use of antibiotics drove the emergence of multidrug-resistant bacteria (including Methicillin-resistant *Staphylococcus aureus* (MRSA), vancomycin-resistant enterococci and highly-resistant *Enterobacteriaceae*)^1,2^, which poses great challenges to improving clinical cure rates and mandating an effective prevention measure^3–5^.

MRSA, as one of the multidrug-resistant Gram-positive bacteria, is resistant to multiple antibiotic classes. Indeed, *Staphylococcus aureus* can acquire resistance to any antibiotic^6^, which has facilitated the occurrence of accurate and fast antibiotic susceptibility testing (AST) for this pathogen^7–9^. Compared to current gold-standard AST with 48-72 h response^10,11^, newer approaches accomodate a rapid detection of drug-resistant *Staphylococcus aureus* with the advantage of a quicker turnaround time^12–14^. Besides, clinical specimens are able to be directly used for susceptibility testing^15–17^, which provides convenience for sample preprocessing procedure. These methods that depend on molecular detection of gene targets would lead to false negatives^18^. The adoption of matrix-associated laser desorption and ionization/time-of-flight mass spectrometry (MALDI-TOF MS) instruments benefits the rapid pathogen detection within 2 h from subcultured colonies^19^. Many previous works have focused on the discrimination between MRSA and methicillin-susceptible *Staphylococcus aureus* (MSSA), which requires the identification of spectral peaks for MRSA and MSSA^20–22^. However, the investigation of MS spectra for other antibiotics to predicting susceptibility is clinically necessary to direct prescription and patient care.

Machine-learning-based (ML-based) techniques have facilitated the analysis of large-scale data from clinical cases. The abundance of MS spectra for clinical specimens collected from *Staphylococcus aureus* infected patients are key for the development of predictive models to assist in diagnosing MRSA. However, single model for the methicillin or oxacillin resistance prediction is not satisfactory for multi-drug resistant *Staphylococcus aureus*. Moreover, models for the susceptibility of other antibiotics are worthy for effective therapies. Further works are needed to achieve a more comprehensive assessment of a multi-drug resistant clinical case than a single diagnosis.

In this work, we develop a XGBoost system to predict whether a *Staphylococcus aureus* infected patient would carry multi-drug resistant *Staphylococcus aureus* on the basis of a 7-year longitudinal study, over 20,000 individually AST results and state-of-the-art machine learning methods. First, any clinical specimen that contains *Staphylococcus aureus* was allowed to conduct further experiments for obtaining MS spectra and AST results on six antibiotics. Second, analyzing the six drug resistant ratios for all samples, we find extremely disproportional rate for penicillin and fusidic acid and strong drug-specific association with resistance (Supplementary Table. 1 and 2). Because almost all isolates were resistant to penicillin and susceptible to fusidic acid, these two antibiotics were excluded from the construction of predictive models. Third, instead of a single classification between MRSA and MSSA, we construct four predictive models, that are of differing features and parameters and guide clinicians to treat infected patients within the next 24 h. Finally, patient-specific drug usage could also be reduced efficiently by these models, thereby alleviating the burden for both patients and health care community. The machine-learning-guided personalized empirical prescription will minimize medication failure and reduce the overall use of antibiotics in the long run, applied in the clinic, thereby aiding in the worldwide campaign to impede the epidemic of antibiotic resistance.

## Results

### Data and cohort characteristics

An overview of the cohorts is described (Fig. 1). We included a total of 29685 patients between 2013 and 2019 in Linkou cohort (Fig. 1a; see Methods). 215 patients were excluded because their clinical specimens were notified of negative screening results for *Staphylococcus aureus*. 26852 patients (after exclusion of an additional 83 patients with missing covariates and 2535 patients with missing AST results). For external validation, 5303 patients were assigned to the replication population from Kaohsiung cohort (Fig. 1a; see Methods). With the same experimental workflow, an independent test set that included 4955 patients (after exclusion of 76 *Staphylococcus aureus* negative specimens, 3 patients with missing covariates and 269 patients with missing AST results) was used for external validation. Key patient and clinical specimen characteristics are shown in Fig. 1b, c. In both Linkou and Kaohsiung cohort, nearly 60% of infections were male. Any sample type was allowed in the experimental design and wound is probably the most commonly processed form of *Staphylococcus aureus* positive clinical sample in the body (Fig. 1b). The age density plot (Fig. 1c) explicitly focuses on the display of age density distributions of the two cohorts. The peaks of both density plots display that the ages are concentrated over the interval (60, 70) and near-0 which indicates the more susceptible population of *Staphylococcus aureus*.

**Fig. 1.**
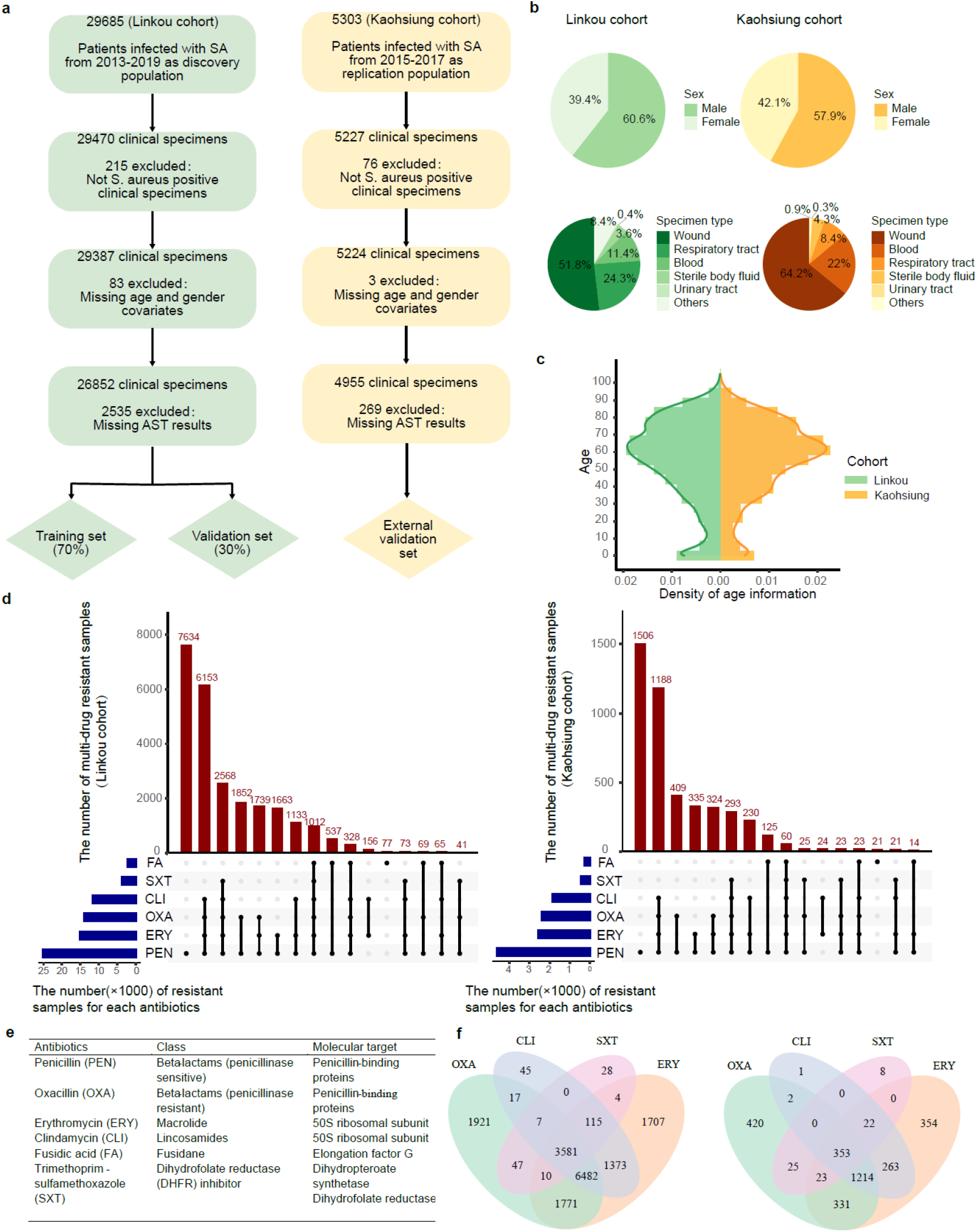
**a-c, Data and cohort characteristics. a,** Cohort selection. *Staphylococcus aureus* positive specimens were first screened. Next, patients with missing age, gender covariates and AST results were excluded. Finally, the cohort was divided into training and validation sets (see Methods). **b**, Basic characteristics of the cohort data. Pie charts are divided according to the sum of data points in sex and specimen types. **c**, Distribution of ages for both cohorts, respectively. **d-f, Multidrug-resistant *Staphylococcus aureus* isolates in the two cohorts. d,** blue horizontal bar, the number of samples that is susceptible to PEN, ERY, OXA, CLI, SXT and FA, respectively. red vertical bar, the number of samples that is non-susceptible to different combinations of six antibiotics listed in Fig. 1e. **e**, List of antibiotics analyzed in the study. **f**, Venn diagram of the number of resistant samples under the four conditions, excluding PEN and FA. PEN, Penicillin; ERY, Erythromycin; OXA, Oxacillin; CLI, Clindamycin; SXT, trimethoprim-sulfamethoxazole; FA, Fusidic Acid.

### Multidrug-resistant *Staphylococcus aureus*

The study primarily focusses on resistance to the six drugs that were most commonly prescribed as part of the empirical treatment of *Staphylococcus aureus* infection (Supplementary Table. 1–2; Methods). Fig. 1e shows the class and molecular target for the six antibiotics used in this study. Antibiotic categories and the target sites of antibiotics are various. The isolate that is non-susceptible to different combinations of six antibiotics is counted for Fig. 1d. In both cohorts, the top two conditions were that the isolate was only resistant to penicillin and non-susceptible to 4 antibiotics (PEN, OXA, CLI and ERY) agents. Moreover, it must be emphasized that large-scale drug resistance comparing susceptible cases with sensitive controls have identified multi-drug resistance isolates associated with two cohorts, suggesting that multi-drug resistance in *Staphylococcus aureus* has become a very common phenomenon that is a potential obstacle for improving clinical treatment effectiveness (Fig. 1d, f). While, individually, the associated antibiotic has been shown with different proportions of resistant samples in both cohorts (Supplementary Table. 1). For all six drugs, an extremely higher resistance rate in Linkou cohort (93.7%) and Kaohsiung cohort (93.1%) was strongly associated with penicillin. On the contrary, the resistant rate was the lowest for fusidic acid in Linkou cohort (8.3%) and Kaohsiung cohort (5.4%). Hence, the other four drugs become the focus of our research attention. However, limited information exists regarding the relationship of drug-resistant *Staphylococcus aureus* over the four antibiotics.

### Preparation of a high resolution MALDI-TOF mass spectra dataset

A summary of our workflow is demonstrated in Fig. 2. Generation for MALDI-TOF MS spectra of all the samples described in the Method section is shown in Fig. 2a. All spectra are shown scaled peak intensities in the Y-direction, covering a range of 2,000–20,000 Dalton (Da) in the X-direction. The raw MS spectra were converted into peak lists (Methods), which were then used for model development after data pre-processing by binning normalization (Fig. 2b; Methods). The features generated for the prediction task include 900 intensities of pseudo-ions for each sample.

**Fig. 2.**
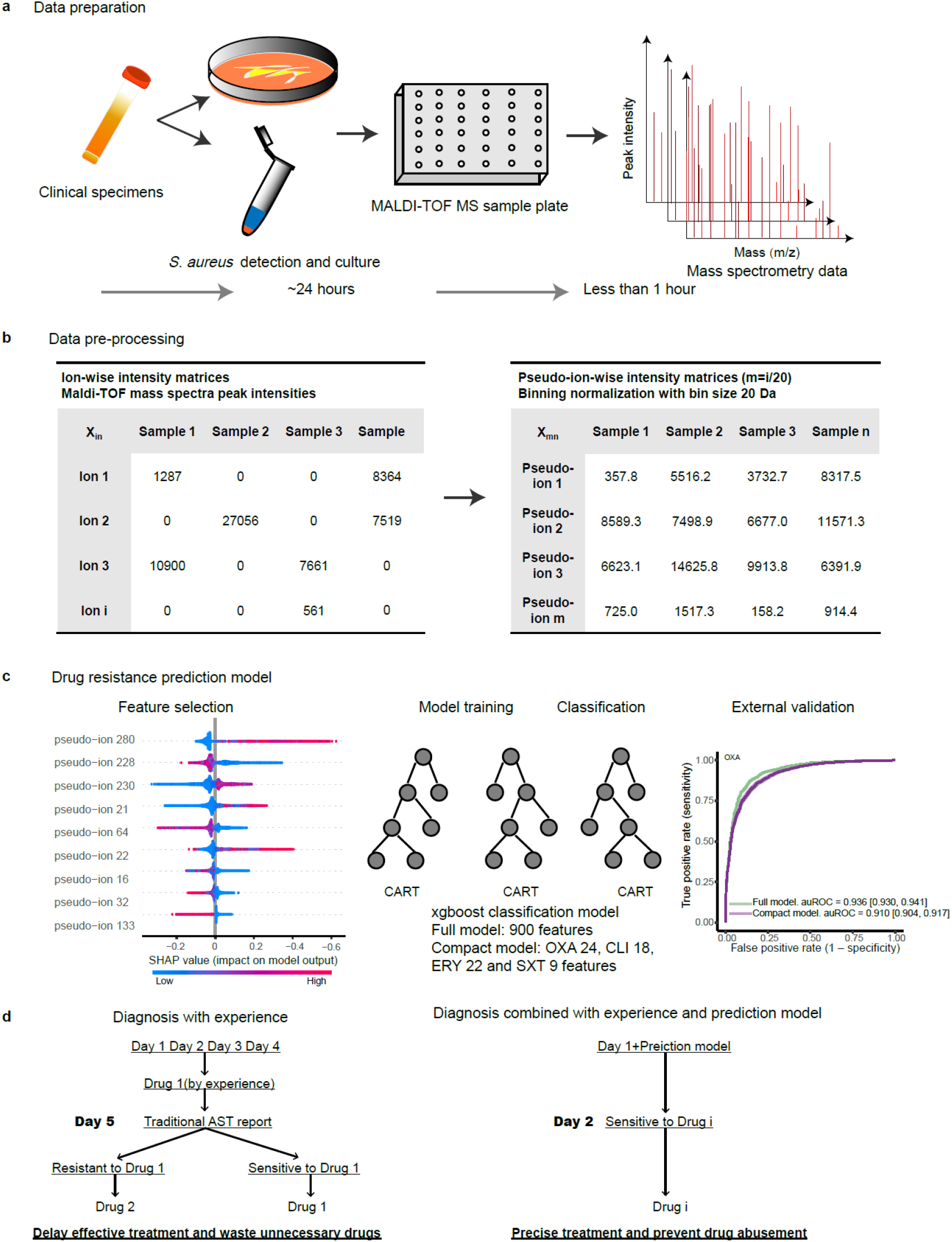
Model development overview. **a**, Data preparation. Clinical specimens were exported from hospitals to microbiology laboratory and AST was conducted on the cultured samples by conventional methods. At the same time, the generation of MALDI-TOF MS spectra for each sample was ensured. **b**, Data pre-processing. MS data normalization was performed by intensity norm averaging algorithm and the m/z in a 20 Da grid (method). **c**, Drug-resistance prediction model. All features, together with drug resistance information representing the susceptibilities of all the samples for each drug, were used to construct the full models for the binary classification problem. The features were annotated according to the absolute mean SHAP value and four feature sets formed the four compact models. The classifiers were trained on the discovery population by the four feature label sets. An XGBoost model was chosen as the classifier which was evaluated on the replication population as the external validation. **d**, Clinical drug recommendations with experience and prediction model. AST, antibiotic susceptibility testing; MS, mass spectrometry; m/z, mass to charge ratio; Da, Dalton.

### Development of models for prediction on multi-drug resistance

We aimed to determine the multi-drug resistance of a patient developing *Staphylococcus aureus* infection within 24 hours using ML-based mass spectrometry analysis. Owing to the disproportional resistant samples for penicillin and fusicid acid, the susceptible labels generated for the prediction task included oxacillin, clindamycin, erythromycin and trimethoprim-sulfamethoxazole. The analysis framework for the feature selection and model building was shown in Fig.2c. For all the samples of the discovery population as the training set, the dataset contained 900 intensities of pseudo-ions as features and four label sets obtained by drug susceptibility testing of *Staphylococcus aureus* based on the four antimicrobial agents. Four XGBoost classifiers were built up on the discovery population and SHAP (SHapley Additive exPlanations) value-based features formed the compact models. Two XGBoost classifiers with each drug were developed—the full and compact models. There are 900 features generated from the MS spectra data of the discovery population that were ranked according to the mean absolute SHAP value^23^, which indicates their contributions to the prediction models. The full models used 900 features, originating from 18000 ion peaks and the compact models used 24, 18, 22 and 9 features (Supplementary Fig. 1). In clinical practice, the model impact on the prescription was evaluated, based on the 1-year test trail on the usage for drugs (Fig. 2d).

### Inspection of model features

The features generated for the prediction tasks were selected and assessed by mean absolute SHAP values (see Methods). In Fig. 3a, we list the common features shared by four compact models and show the heatmap of SHAP values at m/z ranging from 2,000 to 10,000. We observed that the SHAP value files of five common pseudo-ions from the models were similar in the relevant m/z intervals than other ranges, reflecting the very similar patterns released from different antibiotics. Moreover, analyses of other features of different drug models (Fig. 3b) showed that some features appeared unique to some drugs, indicating the potential different diffusion characteristics of these antibiotics. Summary plots of the entire feature set for each of the compact models related to different investigated bacterial-resistance classifications demonstrates the relationship between features’ original values and corresponding importance (Fig.4c, Supplementary Fig. 1c, 2c, 3c). Furthermore, to capture the association of drug resistance risk and a specific feature, dependence plots were built in the form of drug resistance risk against the pseudo-ion’s peak intensity (Fig.4d, Supplementary Fig. 1d, 2d, 3d; see Methods). The relationship between feature value and SHAP value is illustrated for the top-ranked features of four drug models. Of note, pseudo-ion 21 (m/z range 2401-2420) and pseudo-ion 230 (m/z range 6581-6600) could be a risk factor that contributes to drug resistance. On the contrary, higher peak intensities shown at pseudo-ion 228 (m/z range 6541-6560) decrease the risk for drug resistance. As expected, the drug resistance risk increases as the intensities of these m/z intervals (2441-2460, 3021-3040) increase^8,21^.

**Fig. 3.**
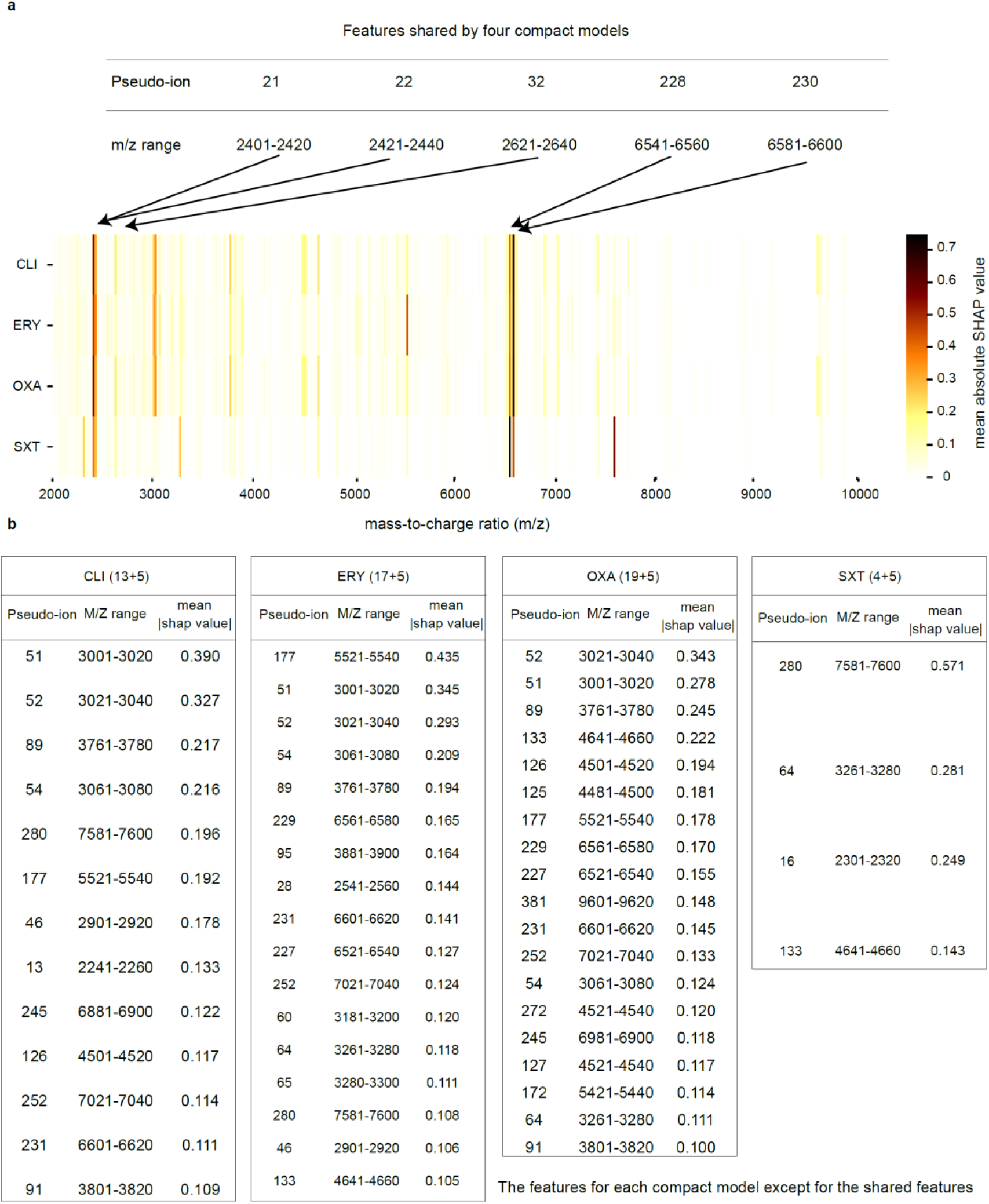
Inspection of model features. **a,** Heatmap of mean absolute SHAP value of features shared by four compact models. For better visualization, features were converted from pseudo-ions to their corresponding m/z values. **b**, Pseudo-ions, m/z ranges and mean absolute SHAP values were summarized for each feature set. SHAP, SHapley Additive exPlanations.

**Fig. 4.**
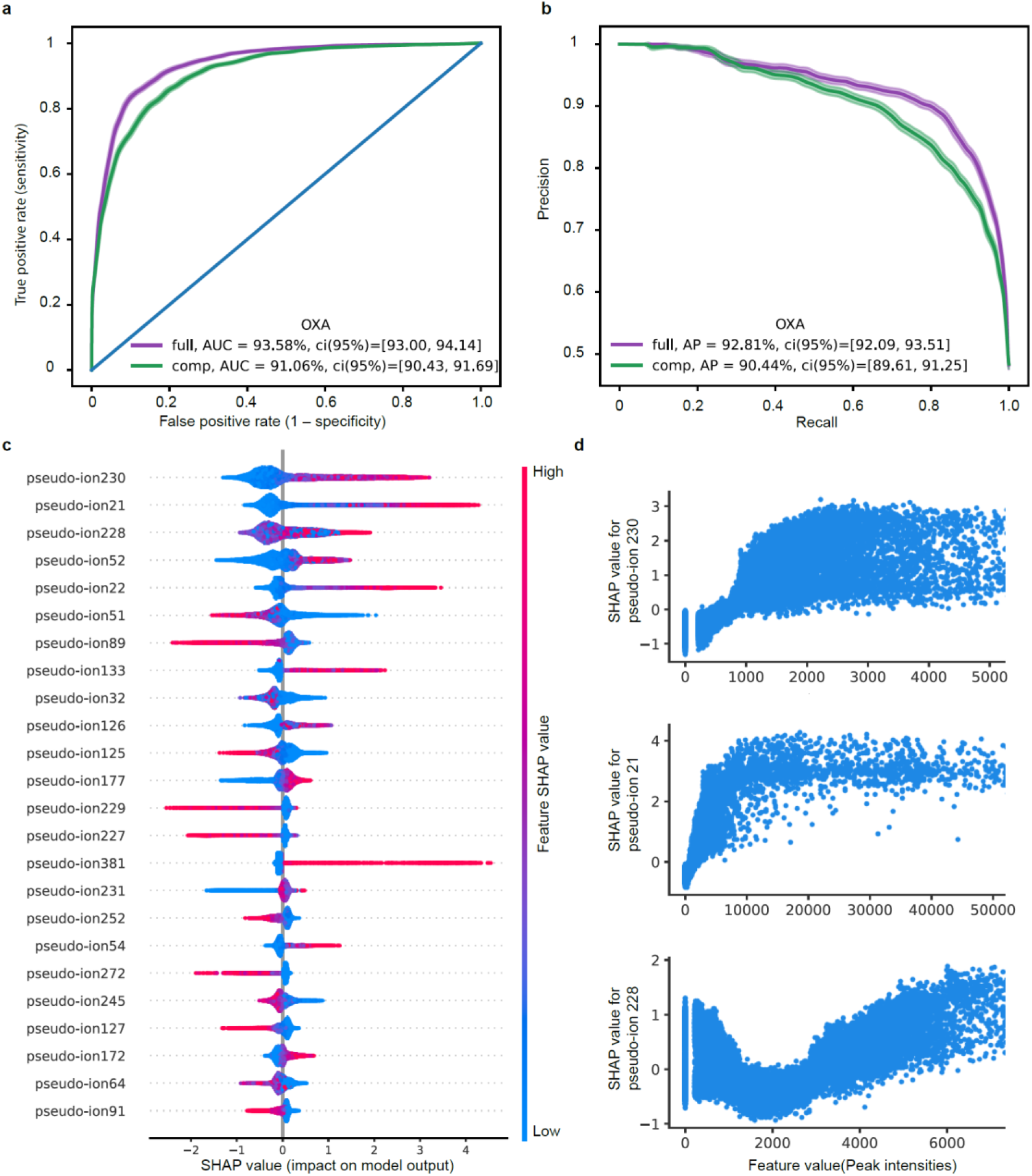
Model features and performance. **a**, Receiver-operating characteristic curve of the binary classification task of OXA resistance, comparing the full model with the compact model. The full model contains 900 features (pseudo-ions), and the compact model contains 24 features. **b**, Precision-recall curve for the full and compact classification model. **c**, A summary plot of the SHAP values for the feature set of the compact model. Features are ranked by their overall importance in creating the final prediction. For each feature, every point is a specific sample, with colors ranging from red (high values of the predictor) to blue (low values of the predictor). **d**, Dependence plots for the top 3 features with the largest mean absolute SHAP value, showing peak intensities versus its SHAP value in the prediction model. OXA, oxacillin; CLI, Clindamycin; ERY, Erythromycin; SXT, Trimethoprim-sulfamethoxazole; comp, compact; AUC, area under the ROC Curve; AP, area under the precision recall curve.

### External validation

The replication population of Kaohsiung cohort was used for external validation. We performed identical data pre-processing strategies as the discovery population of Linkou cohort. Models were evaluated on the performance of drug resistance prediction for the same four antibiotics. The testing results report that the performance of the areas under the receiver operator curves (AUROCs) of the full and compact models were 0.936 and 0.911, 0.896 and 0.876, 0.860 and 0.842, 0.908 and 0.892 for OXA, CLI, ERY, SXT, respectively (Fig. 4a, Supplementary Fig. 1a, 2a, 3a). Additionally, for each antibiotic, the performance of our full and compact models is shown in Fig. 4b (Supplementary Fig. 1b, 2b, 3b), using precision-recall curves^24^ (PRC). The areas under the precision-recall curves (AUPRCs) are more informative and were 0.928 and 0.904 (OXA), 0.866 and 0.841 (CLI), 0.873 and 0.859 (ERY), 0.814 and 0.795 (SXT) for the full and compact models, respectively. We observed a slight performance decrease for all the compact models which obtained result in the accurate AST classification and valuable characterization of features, including intensity of m/z of unexplained resistance. Considering the imbalanced dataset for SXT (Supplementary Table. 2, 14.1%resistant samples and 85.9% susceptible samples for Linkou cohort; 8.6% resistant samples and 91.4% susceptible samples for Kaohsiung cohort.), as expected, the recall rates of the SXT models show a slightly lower performance compared with the other models from a more balanced dataset.

### ML-based drug recommendations substantially reduce the usage of unnecessary drugs

Across the 1-year test period starting from July 2018 to June 2019, cephalosporins usage for 860 patients were recorded (Supplementary Table. 3). Totally 2516 inappropriate cephalosporins were avoided for treating MRSA. 743.4 daily defined dose (DDD) of cephalosporins were reduced by the oxacillin resistant prediction model. In fact, all cephalosporins except fifth generation cephalosporins are not considered efficient in treating MRSA. In the clinical scenario, misusing third generation cephalosporins for treating MRSA would be fairly common before the final ASTs showing that the pathogens are actually MRSA. It is in concordance with the fact that the majority of the reduced cephalosporin was third generation cephalosporins (Supplementary Table. 3). In addition to third generation cephalosporins, totally 49.8 DDD/398 doses of fourth generation cephalosporin (i.e. Cefepime) could be avoided by using the ML-based drug recommendation.

## Discussion

Our study provides a rapid susceptibility testing pipeline for the early detection of the multi-drug resistant *Staphylococcus aureus* of effected individuals. By integrating MALDI-TOF MS spectra information and ML methods, this work offers several advantages over the current gold standard AST assays. First, as with other AST assays, our pipeline determines the susceptibility to many antibiotic agents of *Staphylococcus aureus* infected patients, a clear advantage over single drug AST assay. Second, our AST results could be obtained within 24 h from the time of bacterial culture, compared with 24-72 h by the conventional methods. Third, the machine learning approach to binary classification provides actionable AST indication that can assist clinicians of reasonable antibiotic choice. Fourth, the implications of the model features give us a better insight into the common or special characteristics of different antibiotic classes. Fifth, ML-based drug recommendation could effectively help to reduce the unnecessary drug usage and guide the physician to prescribe medication.

Our experimental design allows a wide range of clinical samples, and almost any *Staphylococcus aureus* infected specimen types (e.g. wound, respiratory tract, blood, sterile body fluid, and urinary tract) are accepted for further subculturing and generating MS spectra. Additionally, we built and replicated binary classification models for the prediction of multi-drug resistance in patients with *Staphylococcus aureus* infection that accurately forecast oxacillin-resistance, erythromycin-resistance, clindamycin-resistance, and trimethoprim-sulfamethoxazole-resistance with an area under the curve of 0.94, 0.90, 0.86 and 0.91 in the replication population, respectively.

Built starting from a full set of features from MS spectra collected from longitudinal data cohort, the predictive algorithm developed has strengths. Long-term follow-up data was able to fully utilize the data information for all the full models. Then the features were winnowed down through the SHAP value. Only a few features that contributed the most formed the compact models. Although the AUC for compact models decreased slightly compared with the full models, such classifier performance was clinically acceptable and features may provide an independent valuable characterization of the resistance mechanism. MALDI-TOF incorporated with the ML-based clinical diagnostics could help clinicians to accurately prescribe drugs and reduce the misusage of unnecessary drugs.

The main limitations of our study are related to the single-area design, which is constrained within the Taiwan area. However, the recruitment patients of the longitudinal study are from two hospital centers (Linkou and Kaohsiung), covering more than 20,000 patients from 2013 to 2019. Such large-scale validation demonstrates the applicability and reproducibility in other areas. Furthermore, the investigated features were not satisfied to conclude the potential gene or proteins that are responsible for the drug resistance because only ions were obtained from the MALDI-TOF MS spectra based on our experiment design. Further development will be needed to detect specific genes or proteins by MS/MS spectra according to the promising m/z intervals.

In conclusion, here we demonstrate a comprehensive analysis framework including data pre-processing, feature selection and interpretation, and XGBoost hyperparameter tuning and model building to construct the rapid platform for AST that leverages the advantages of both MALDI-TOF and machine learning. Considering the verified good performance of our models, we postulate that MALDI-TOF and ML-based AST method may help clinicians to accurately make decisions to prescribe drugs. Although a substantial reduction in the usage of impropriate antibiotics, it is agnostic to the mechanism of resistance. Nevertheless, it is a first step for the rapid AST for multiple drugs in the clinical studies that investigate the potential features of early interventions for antibiotic resistance. Equally, these ML-guided AST approaches can further implement in the clinic and be extendable to any pathogen and antibiotic class, thereby reducing unnecessary medical expense and impeding the drug resistance epidemic.

## Methods

### Ethical approval and patient consent

This was a retrospective study investigating the relation between MS spectrum and microbial strain typing. No diagnosis or treatment was involved in the study. Waiver of informed consent was approved by the Institutional Review Board of Chang Gung Medical Foundation (No. 202100008B1).

### Hospital Cohort

The data that was consecutively collected from Linkou Chang Gung Memorial Hospital for the period 2013-2019 that were assigned to discovery population (Linkou cohort). Independent replication (Kaohsiung cohort) was attained in patients from Kaohsiung Chang Gung Memorial Hospital cohort for the period 2015-2017. Microbiology culture and antibiotic susceptible test were conducted in the CGMH clinical microbiology laboratory. From the hospital cohort, we identified a cohort of patients who infected with *Staphylococcus aureus* and kept the positive samples for further AST.

### Trial design and strain acquisition

For the development and validation of a clinical prediction model, the study was planned as a retrospective cohort study. We conducted the trial as a two-cohort (two medical centers) to testify the effect of the antibiotic susceptible test with mass spectrometry on the multi-drug resistant *staphylococcus aureus*. All bacterial cultures in the two medical centers were tested in the clinical microbiology laboratories. Depend on the location of the suspected infection, a sample was taken from wound, respiratory tract, blood, urine, sterile body fluid, or other parts of body. Clinical specimens that tested positive for *staphylococcus aureus* were kept. The distribution of the origins of the specimens is summarized in Fig. 1b. Same cultured bacterial isolates were used for both AST and MALDI. Mass spectrometry data were obtained with the use of MALDI-TOF mass spectrometry, and antibiotic susceptible test was undertaken by disk diffusion method and broth microdilution, that revealed susceptibility or resistance to penicillin, oxacillin, erythromycin, trimethoprim-sulfamethoxazole and fusidic acid, respectively.

#### MALDI-TOF mass spectra experimental conditions

Mass spectrometry was carried out using a Bruker Microflex LT MALDI-TOF system (Bruker Daltonik, Bremen, Germany) as described previously^8^. The operation of the Microflex LT was followed by the manufacturer’s instructions. Fresh cultured isolates were smeared to the MALDI steel 96-well target plate by the operator. Extraction with formic acid (1 μL, 70%) was performed on a thin film. After drying at 25 °C, the target plate was covered using matrix solution that comprises a mixture of solvents (1%α-cyano-4-hydroxycinnamic acid in 50% acetonitrile containing 2.5% trifluoroacetic acid). When the samples were dried at room temperature, the target plate was loaded in to the analyzer and it was analyzed by microflex LT MALDI-TOF analyzer operated in linear ionization mode (accelerating voltage, 20kV; nitrogen laser frequency: 60 Hz; 240 laser shots). The spectra were recorded with the mass/charge ratio (m/z) ranging from m/z 2,000 to 20,000. Peak patterns were analyzed in R using the R package MALDIquant^25^. The raw spectra were undergoing a three-step process. Firstly, baseline correction (Top-hat filter) was applied. Then the peaks were determined by calculating the median absolute deviation (MAD) and the half window size was 10. Finally, peaks with signal to noise ratio (SNR) ≥ 5 were collected for the upcoming analysis. After the further processing of the raw MS spectra, a peak list containing m/z values and intensities, the so-called ‘mass fingerprint’ of a sample is developed.

#### Data Preprocessing

‘Mass fingerprint’ of a sample consists of the ion mass-to-charge ratio (m/z) and raw intensity values of all peaks. Preprocessing steps were applied to translate original ion intensities to relative pseudo-ion abundance. This was a three-step process and was modified from a previously described method^8^. First, ion peaks with an intensity <= 100 were removed from the analysis. Second, for each sample with its m/z ranged from 2,000 to 20,000, m/z axis was split into equal intervals with bin size 20^8,21^. (20,000-2,000)/20=900 vectors were obtained and named as pseudo-ions for further analysis. Third, the normalization within each interval vector was applied. This was achieved by calculating the *l*_1_-norm of the interval vector divided by its *l*_0_-norm and then added its *l*_2_-norm, obtaining the normalized intensity for the pseudo-ion.

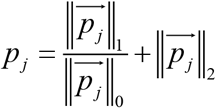

Where 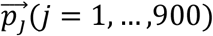 corresponds to the 900 interval vectors, *p_j_*(*j* = 1, …,900) is the normalized intensities of the 900 pseudo-ions. ∥•∥_0_, ∥•∥_1_ and ∥•∥_2_ represent the *l*_0_-norm, *l*_1_-norm and *l*_2_-norm respectively. Last, every sample followed the same steps to obtain their normalized pseudo-ion intensities. A pseudo-ion matrix table was then produced, which included all cohorts and their corresponding preprocessed mass spectrometry data across the whole study. For each antibiotic, a drug-resistant or drug-susceptible group is defined as samples that are identified by the susceptibility result (i.e., those that were labeled as susceptible or resistant).

### Feature definition and selection

Every mass spectrum is a plot of peak intensities of the mass-to-charge ratio. However, to apply machine learning methods, a fixed-length feature representation is necessary. Following the data preprocessing step, with bin size 20 as equal intervals, 900 intensities of pseudo-ions were generated for each m/z interval. If no peak occurrence in some m/z intervals, the corresponding intensities would be zero. These 900 features were provided to the full models for the four-drug resistance prediction classifiers. The importance of individual features was measured using mean absolute SHAP values made on the dataset for the discovery population. In each model, implemented for all antibiotics, the selected features for the compact model were obtained with the settled threshold by which the mean absolute SHAP value larger than 0.1. The resulting 24, 18, 22 and 9 features, along with AST susceptibility classification for each training set, formed the corresponding compact models.

#### XGBoost classifier

Classifiers for prediction were generated using an XGBoost model^26^ built with the tree booster. XGBoost is an efficient implementation of a gradient boosting framework for handling sparse data. We used the Python module xgboost to infer parameters for a classifier given labels of the resistant or susceptible group. Booster was set to gbtree, objective was set to binary: logistic, and the evaluation matric was set to AUC. In both discovery population and replication population, the ratios of penicillin- resistant samples and fusidic acid susceptible samples are extremely high (**Supplementary Table**. 2). Therefore we excluded drug-resistant prediction for penicillin and fusidic acid. For the other four antibiotics, the training thus comprised four paralleled processes with four settings in which the models may be implemented in practice.

XGBoost classifier depends on its optimized parameters, including the number of iteration (nrounds), the learning rate (eta), and the maximum depth of a tree (max_depth). Grid search approach was with the following search range: nrouncds ∈ [40-200, with an interval of 10], eta ∈ [0.0001, 0.001, 0.01, 0.05, 0.1, 0.2, 0.25, 0.3, 0.5], max_depth ∈ [2, 4, 6, 8, 10, 12]. Hyperparameters were selected following a cross-validated grid search, with the following settings selected (Supplementary Table. 4).

XGBoost classifiers were first trained on a subset of the discovery population of Linkou cohort (that is, the training set), and then were applied to the remaining data (that is, validation set), to infer the ability of the classifier to classify new data. A bootstrap sample size was set to 70% of the training set. The purpose of CV is not to build models but to assess the stability of the model performance. Final models were built on the whole discovery population based on the trained hyperparameters.

### External validation on Kaohsiung cohort

To evaluate the predictive ability of the classification scheme while considering an independent patient cohort. The replication population of Kaohsiung cohort was used for external validation. We performed identical data preparation and pre-processing with the discovery population from Linkou cohort. Since the disproportional resistant samples of the drugs penicillin and fusidic acid, we applied the trained XGBoost classifier models on the Kaohsiung data. To predict the drug resistance for Oxacillin, Erythromycin, Clindamycin, trimethoprim-sulfamethoxazole. We used standardized classifier parameters and standardized thresholds that were inferred based on the same training set to generate a series of classification models on each drug resistance labels and assessed the accuracy and discriminatory power of these models using AUROC and AUPRC. For a developed classifier, the assessment was based on its accuracy and discriminatory power. A receiver operating characteristic curve, or ROC curve, illustrates the diagnostic/classification ability of a binary classifier system as its prediction threshold is varied from 1 to 0. True positive rate (sensitivity) and false positive rate (1-specificity) formed the ROC curve. Whereas AUROC varies between 0 and 1 — with a random choice yielding 0.5, to 1.0, an excellent classifier.

### Model interpretations

To discovery the feature importance and the relationship of individual features with the models, SHAP values^27^ were used to evaluate the feature contribution for model performance. How the impact of variables taking into consideration interacted with other variables were measured by SHAP values. For each sample, the predicted result was separated by SHAP values into the contribution of every feature value. In this study, the Shapley value is used to discover the feature importance and evaluate how each feature makes impact to the complicated model. It originates in game theory approach, can segregate the prediction outcome of each sample to the constitution of the feature contributions.

Denote *x* as the input pseudo-ion vector for a sample. The *f*(*x*) is the predicted outcome by the classifier *f*(·). The Shapley analysis can be given by the following equation:

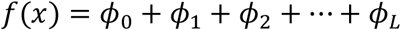

where *ϕ*_0_ = *E*[*f*(*x*)] is the base Shapley value generated by the expectation of the model output over the training set. The *ϕ_l_* = *ϕ_l_*(*f, x*), *l* ∈ {1,2, ···, *L*} are the Shapley values related to the *L* pseudo-ion features, which can represent the impact of the feature *l* to the predict outcome *f*(*x*).

### Clinical analysis of drug usage based on the prediction model

To evaluate the clinical impact on the AST prediction model, we recorded the antibiotic usage for 860 cases across the 1-year test period (2018.7-2019.6). Oxacillin-resistant *Staphylococcus aureus* is resistant to most currently available beta-lactam antimicrobial agents, including cephalosporins, which is one of the most common beta-lactam antimicrobial agents. For the patients with *Staphylococcus aureus* infection, cephalosporins were prescribed as a matter of experience and clinicians wouldn’t change drugs until the traditional AST report came out in four days. With our prediction model, once the drug susceptibility was predicted, updated drug recommendations could be set in one day. To quantify and compare the doses of cephalosporins reduced by the algorithmic drug recommendations, we calculated actual total doses of cephalosporins and daily defined dose (DDD) of cephalosporins that is the standardized amount by WHO.

## Acknowledgements

This work was supported by the Warshel Institute for Computational Biology, School of Life and Health Sciences, The Chinese University of Hong Kong, Shenzhen, China and Chang Gung Memorial Hospital (CMRPG3F1721, CMRPG3F1722, CMRPD3I0011).

## Author contributions

Z.W., H.Y.W. and T.Y.L. conceived the project, designed and conducted the analyses, interpreted the results and wrote the manuscript, and are listed in random order. H.Y.W. collected the clinical samples and executed the microbiology experiments. Z.W. conducted the analyses and wrote the manuscript. Y.X.P. assisted with the machine learning analysis and visualization. C.R.C. and J.T.H assisted with manuscript revision. J.J.L. and T.Y.L. supervised the study. All authors have read and approved the manuscript.

---

**Supplementary Fig. 1.**
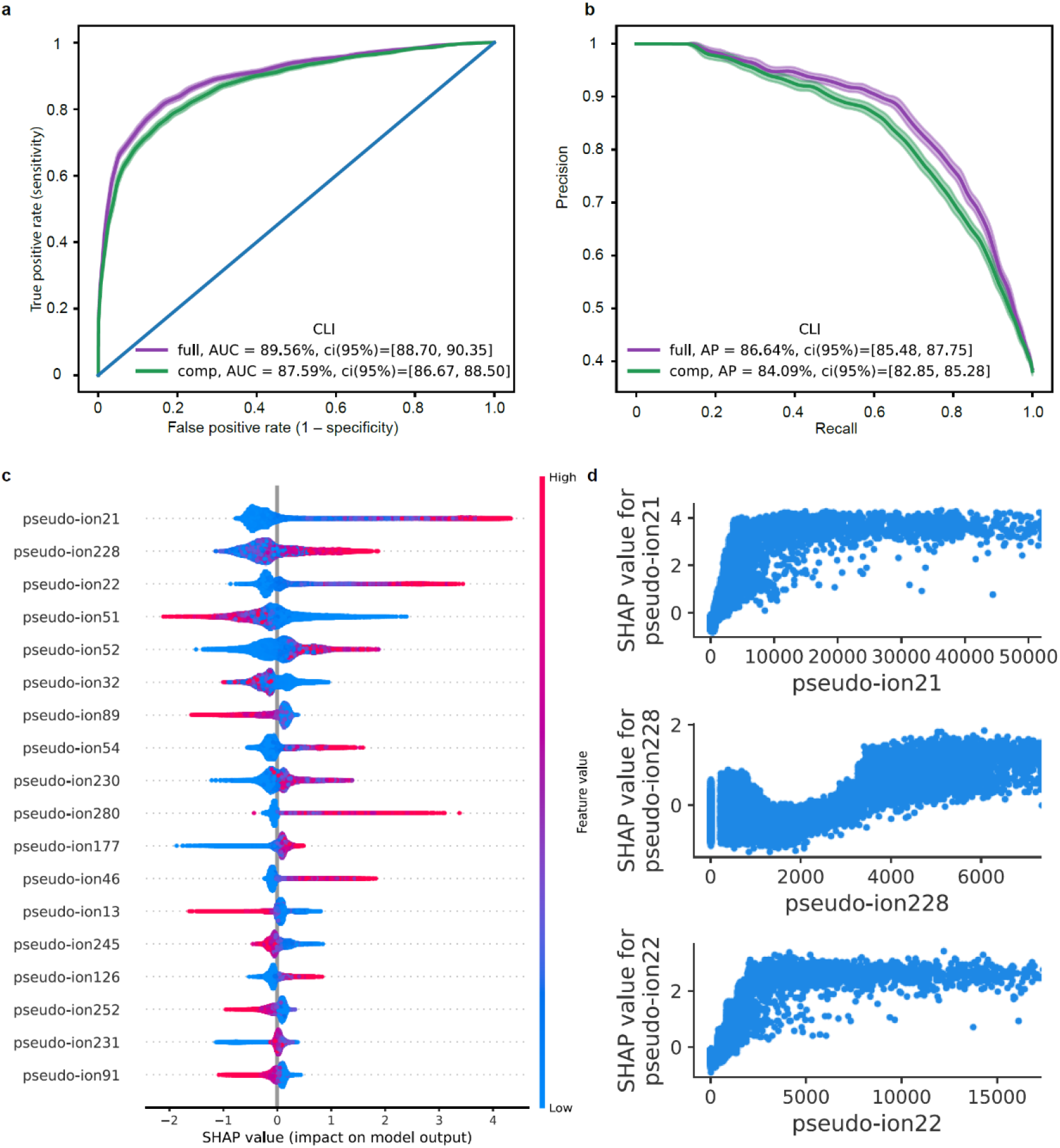
Model features and performance. **a,** Receiver-operating characteristic curve of the binary classification task of CLI resistance, comparing the full model with the compact model. The full model contains 900 features (pseudo-ions), and the compact model contains 18 features. **b**, Precision-recall curve for the full and compact classification model. **c**, A summary plot of the SHAP values for the feature set of the compact model. Features are ranked by their overall importance in creating the final prediction. For each feature, every point is a specific sample, with colors ranging from red (high values of the predictor) to blue (low values of the predictor). **d**, Dependence plots for the top 3 features with the largest mean absolute SHAP value, showing peak intensities versus its SHAP value in the prediction model. OXA, oxacillin; CLI, Clindamycin; ERY, Erythromycin; SXT, Trimethoprim-sulfamethoxazole; comp, compact; AUC, area under the ROC Curve; AP, area under the precision recall curve.

**Supplementary Fig. 2.**
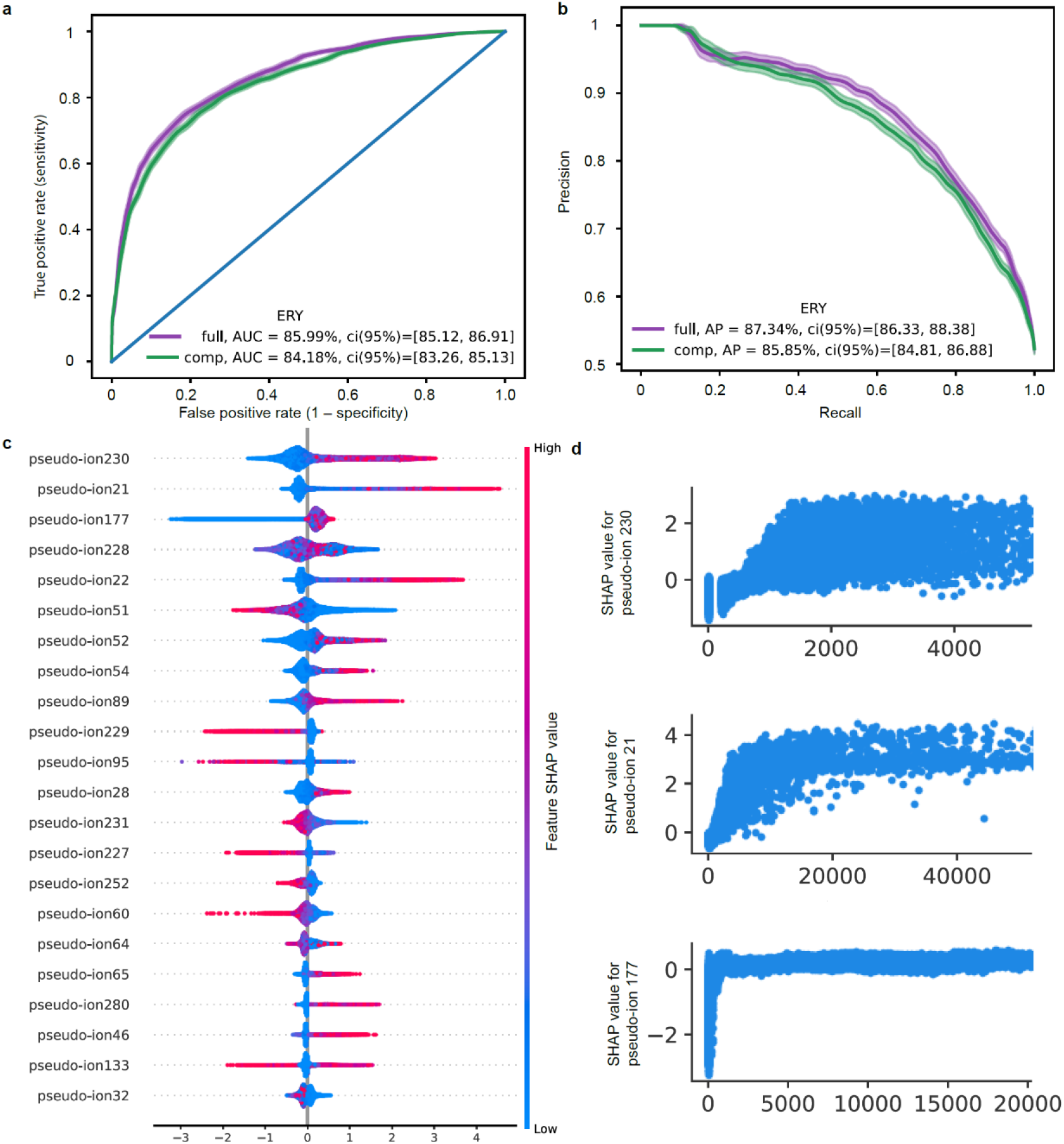
Model features and performance. **a,** Receiver-operating characteristic curve of the binary classification task of ERY resistance, comparing the full model with the compact model. The full model contains 900 features (pseudo-ions), and the compact model contains 22 features. **b**, Precision-recall curve for the full and compact classification model. **c**, A summary plot of the SHAP values for the feature set of the compact model. Features are ranked by their overall importance in creating the final prediction. For each feature, every point is a specific sample, with colors ranging from red (high values of the predictor) to blue (low values of the predictor). **d**, Dependence plots for the top 3 features with the largest mean absolute SHAP value, showing peak intensities versus its SHAP value in the prediction model. ERY, Erythromycin; comp, compact; AUC, area under the ROC Curve; AP, area under the precision recall curve.

**Supplementary Fig. 3.**
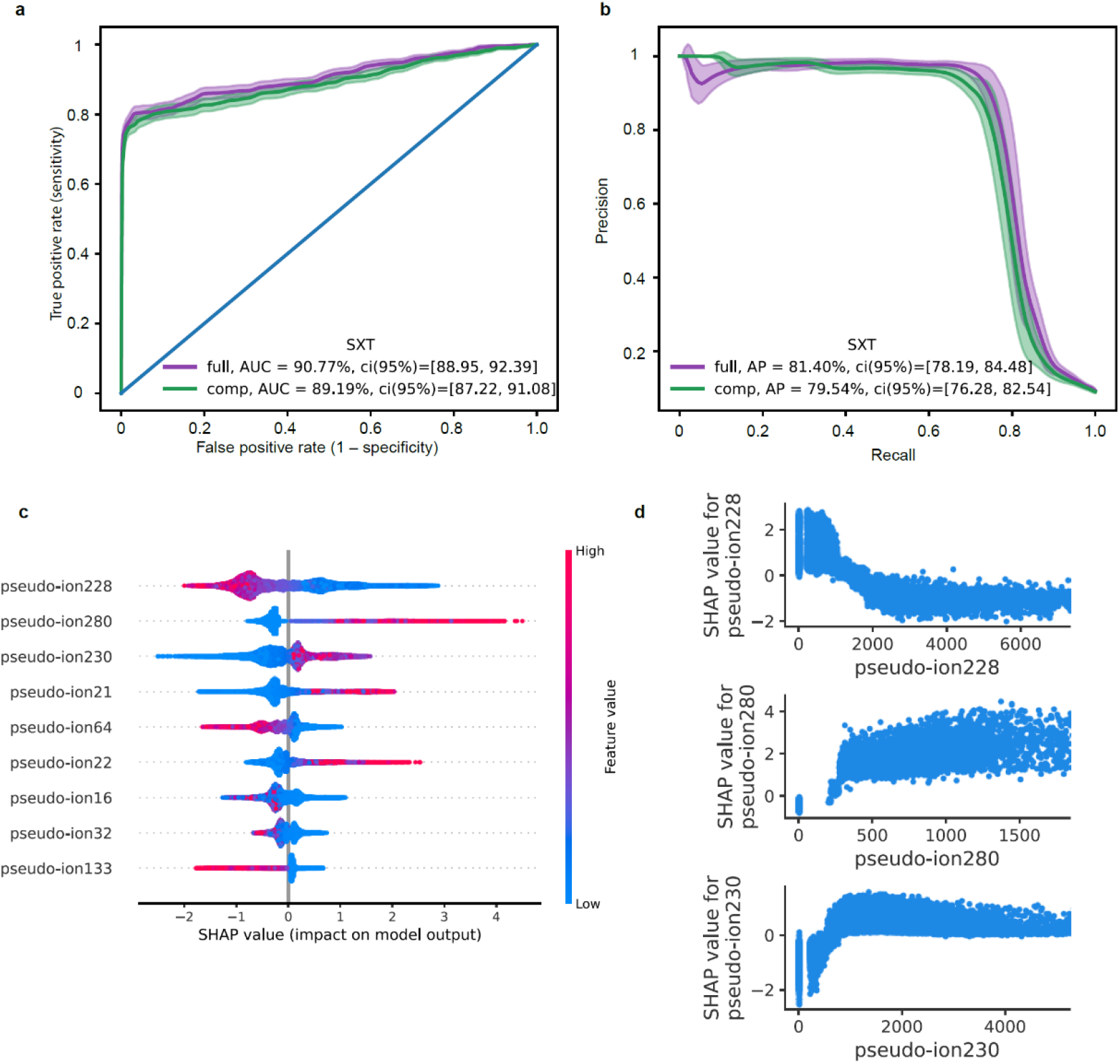
Model features and performance. **a,** Receiver-operating characteristic curve of the binary classification task of SXT resistance, comparing the full model with the compact model. The full model contains 900 features (pseudo-ions), and the compact model contains 9 features. **b**, Precision-recall curve for the full and compact classification model. **c**, A summary plot of the SHAP values for the feature set of the compact model. Features are ranked by their overall importance in creating the final prediction. For each feature, every point is a specific sample, with colors ranging from red (high values of the predictor) to blue (low values of the predictor). **d**, Dependence plots for the top 3 features with the largest mean absolute SHAP value, showing peak intensities versus its SHAP value in the prediction model. SXT, Trimethoprim-sulfamethoxazole; comp, compact; AUC, area under the ROC Curve; AP, area under the precision recall curve.

**Supplementary Table 1.** Drug resistance rate for six antibiotics in two cohorts. Overview of study cohorts. PEN, Penicillin; ERY, Erythromycin; OXA, Oxacillin; CLI, Clindamycin; SXT, trimethoprim-sulfamethoxazole; FA, Fusidic Acid.

| Drugs/Antibiotics | Discovery population<br>(Linkou cohort)<br>26852 samples |  | Replication population<br>(Kaohsiung cohort)<br>4955 samples |  |
| --- | --- | --- | --- | --- |
|  | Resistant<br>samples (%) | Susceptible<br>samples (%) | Resistant<br>samples (%) | Susceptible<br>samples (%) |
| **PEN** | 25151 (93.7) | 1701 (6.3) | 4614 (93.1) | 341 (6.9) |
| **ERY** | 15068 (56.1) | 11784 (43.9) | 2563 (51.7) | 2392 (48.3) |
| **OXA** | 13860 (51.6) | 12992 (48.4) | 2369 (47.8) | 2586 (52.2) |
| **CLI** | 11638 (43.3) | 15214 (56.7) | 1856 (37.4) | 3099 (62.6) |
| **SXT** | 3794 (14.1) | 23058 (85.9) | 431 (8.6) | 4524 (91.4) |
| **FA** | 2233 (8.3) | 24619 (91.7) | 269 (5.4) | 4686 (94.6) |

**Supplementary Table. 2.** Multi-drug resistant proportions in the discovery population of Linkou cohort. PEN, Penicillin; ERY, Erythromycin; OXA, Oxacillin; CLI, Clindamycin; SXT, trimethoprim-sulfamethoxazole; FA, Fusidic Acid.

|  | PEN(%) | ERY(%) | OXA(%) | CLI(%) | SXT(%) | FA(%) |
| --- | --- | --- | --- | --- | --- | --- |
| **PEN** |  | 59.2 | 55.1 | 45.5 | 15.1 | 8.5 |
| **ERY** | 98.8 |  | 78.7 | 76.8 | 24.6 | 10.2 |
| **OXA** | 100.0 | 85.6 |  | 72.9 | 26.3 | 10.5 |
| **CLI** | 98.4 | 99.4 | 86.8 |  | 31.8 | 12.6 |
| **SXT** | 99.8 | 97.8 | 96.1 | 97.7 |  | 28.1 |
| **FA** | 95.6 | 68.8 | 65.0 | 65.7 | 47.7 |  |

**Supplementary Table. 3.** The amount of unnecessary drug usage reduced by the prediction model.

| Cephalosporins | Total Amount | TotalDDD |
| --- | --- | --- |
| CEFAZOLIN SODIUM 1GM/VIAL | 208 | 69.3 |
| CEFUROXIME (SODIUM) 750MG/VIAL | 16 | 4 |
| CEFTRIAXONE 1GM/VIAL | 669 | 334.5 |
| CEFTAZIDIME 1G/VIAL | 547 | 136.8 |
| CEFOPERAZONE SODIUM 500MG +<br>SULBACTAM SODIUM 500MG ) /VIAL ( PC ) | 514 | 128.5 |
| CEFTAZIDIME(TATUMCEF)500MG/VIAL | 164 | 20.5 |
| CEFEPIME 500MG/VIAL | 398 | 49.8 |
| Total | 2516 | 743.4 |

**Supplementary Table. 4.**
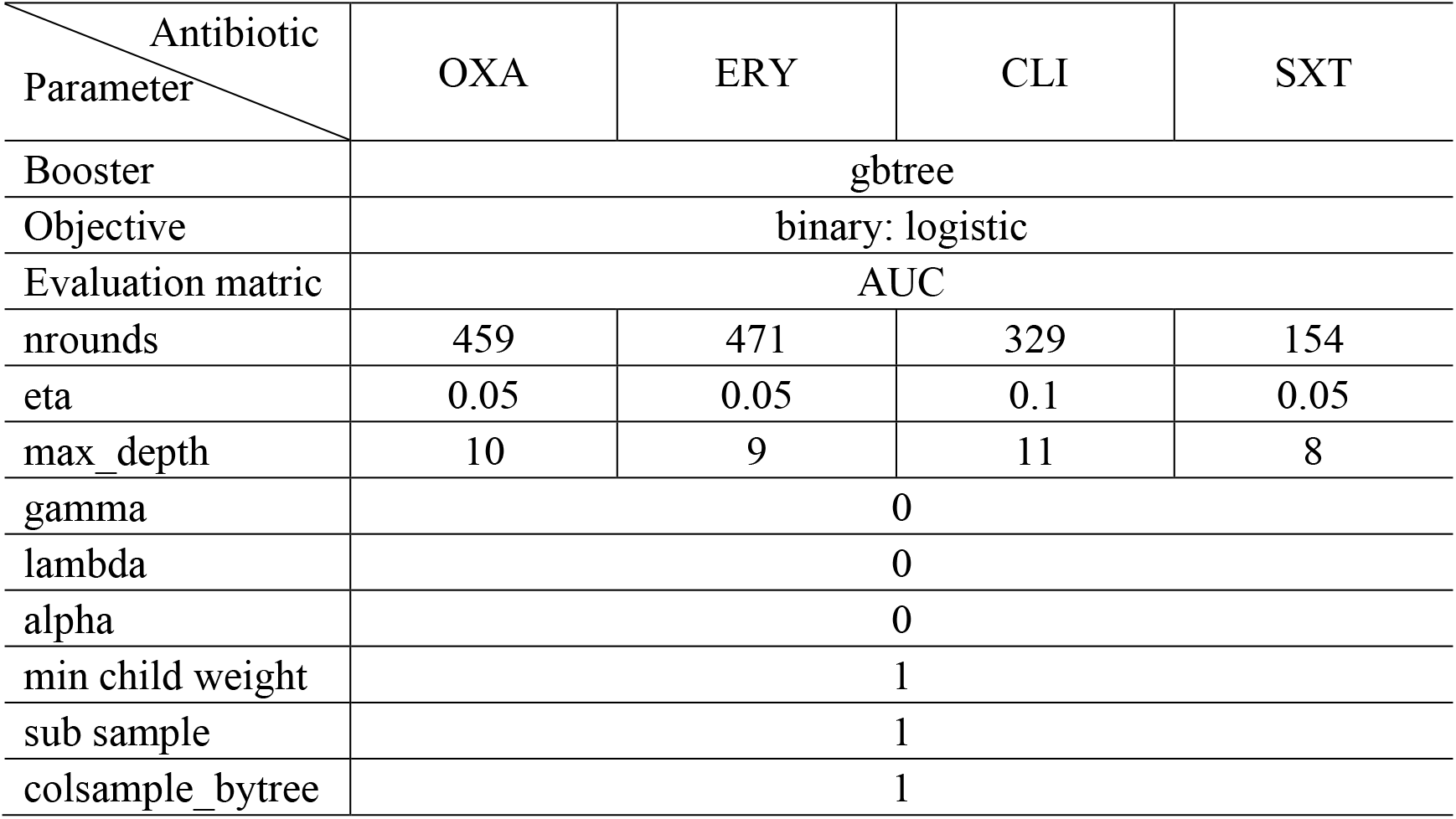
Parameters for python xgboost module.

